# Alignment of multiple metabolomics LC-MS datasets from disparate diseases to reveal fever-associated metabolites

**DOI:** 10.1101/2021.09.02.458540

**Authors:** Ana-Maria Năstase, Michael P. Barrett, Washington B. Cárdenas, Fernanda Bertuccez Cordeiro, Mildred Zambrano, Joyce Andrade, Juan Chang, Mary Regato, Eugenia Carrillo, Laura Botana, Javier Moreno, Kathryn Milne, Philip J. Spence, J. Alexandra Rowe, Simon Rogers

## Abstract

Acute febrile illnesses are still a major cause of mortality and morbidity globally, particularly in low to middle income countries. The aim of this study was to determine any possible metabolic commonalities of patients infected with disparate pathogens that cause fever. Three liquid chromatography-mass spectrometry (LC-MS) datasets investigating the metabolic effects of malaria, leishmaniasis and Zika virus infection were used. The retention time (RT) drift between the datasets was determined using landmarks obtained from the standard reference mixtures generally used in the quality control of the LC-MS experiments. We used fitted Gaussian Process models (GPs) to perform a high level correction of the RT drift between the experiments, followed by standard peakset alignment between the samples with corrected RTs of the three LC-MS datasets. Statistical analysis, annotation and pathway analysis of the integrated peaksets were subsequently performed. Metabolic dysregulation patterns common across the datasets were identified, with kynurenine pathway being the most affected pathway between all three fever-associated datasets.

## Introduction

Many infectious diseases are characterised by fever: a generic host response to numerous microbial pathogens. Fever is associated with the hypothalamus which, in response to the activation of cyclooxygenase-2 (COX-2), releases prostaglandin E2 (PGE2), triggering a systemic increase in body temperature which can have microbicidal effects [1]. Although fever has a protective effect, acute febrile illnesses are still a major cause of mortality and morbidity globally, particularly in low to middle income countries [2]. The failure to correctly diagnose a specific disease associated with fever is partly responsible for this. Inappropriate treatment of misdiagnosed diseases can contribute to the selection of drug resistant microbes. For example, in many parts of Africa, fever is assumed to be due to malaria and treated with anti-malarial drugs. In cases where the patient may actually not have been infected with malaria parasites, but subsequently became infected while the drug concentration was waning, a selective pressure on resistant mutants is imposed [3]. Therefore, improved diagnostics of febrile patients and specific biomarker discovery to support new diagnostics is desirable.

Recently, increasing numbers of studies on fever-associated diseases using untargeted metabolomics with mass spectrometry coupled to high performance liquid chromatography (LC-MS) have emerged [4–8]. LC-MS is a sensitive approach to identifying metabolite markers and for providing a comprehensive coverage of the metabolome, as it enables the separation and measurement of thousands of discrete chemical compounds. Following the LC-MS peak detection process, a list of ions characterised by chromatographic retention time (RT), mass to charge ratio (m/z) and intensity is obtained.

When performed on individual disease states, however, distinguishing between metabolites associated generically with fever, and others specific for particular diseases is a challenge. Thus, we aimed to search for common perturbations to compounds across a set of fever associated diseases simultaneously, in order to identify metabolites generically associated with fever or disease severity. In order to achieve this, a method for integration of multiple LC-MS datasets through peakset alignment was developed.

The alignment process, also referred to as correspondence, for which algorithms are categorized into either direct matching or warping algorithms, has been extensively studied, mainly in the context of multiple injections within the same experiment [9]. This is due to the fact that although the LC-MS instrumentation and methodology are robust and well established, measurement variability can still appear resulting in non-linear shifts especially in RT. There are two types of variability sources which can appear in an LC-MS experiment, systematic and random. Systematic variability is usually consistent and may be caused by factors such as the apparatus itself, the column, system stability and temperature [10]. Random variability, by contrast, has no pattern and cannot be completely removed from a metabolomics experiment although it can be reduced. In order to reduce systematic RT variability within an experiment, a set of known metabolites (standard reference mixture (SRM)) is run at various points during an LC-MS experiment as part of the quality control process [10, 11]. Using SRMs enables drift in compound intensity and RT to be tracked throughout every LC-MS run with SRM metabolites used as landmarks for retention time correction (the m/z for each compound detected using LC-MS being constant).

In this study we used the information provided by SRM runs to determine the RT drift between injections of different LC-MS experiments which was then modelled using Gaussian Process (GP) regression, a supervised machine learning approach [12]. GP regressions are a non-parametric approach to modelling data and they differ from standard regression models in that they do not require any assumptions about a particular parametric form for the function being modelled. We used fitted GPs to perform high level correction of retention times between experiments, after which standard alignment was performed.

The algorithm was developed here specifically to determine whether any metabolites could be identified that changed in abundance in similar ways across a series of distinct fever-associated diseases. These include Zika virus infection in patients from Ecuador [13], Leishmaniasis patients from Spain [14] and uncomplicated malaria infected volunteers from the UK [15]. The samples had all been run previously using the same LC-MS platform (Glasgow Polyomics, University of Glasgow, UK). By seeking metabolites whose variation in abundance followed common trends in different datasets, we aimed to determine disease-generic metabolites that could assist in both understanding the pathophysiology of infectious disease, and also identifying metabolites that change in abundance in individual studies that may be fever rather than specific disease related.

## Materials and methods

### Datasets

Three LC-MS datasets –D_*Z*_, D_*M*_, D_*VL*_– analysed at Glasgow Polyomics metabolomics facility (University of Glasgow, UK) were used for the cross-experimental integration in this study. Detailed information regarding the sample collection process for each dataset is included in the Supporting Information (S1 File).All three experiments were designed for detecting the differences between the serum metabolic profiles of healthy controls and infected patients diagnosed by gold-standard methods. In total, there were 74 samples (37 controls and 37 disease samples). Detailed information about each LC-MS experiment is presented in Table 1.

**Table 1.** LC-MS datasets details.

| | $D_Z$ | $D_M$ | $D_{VL}$ |
| --- | --- | --- | --- |
| Infectious Disease | Zika virus [13] | Malaria [15] | Visceral Leishmaniasis [14] |
| Study Type | Case-control | Intervention | Case-control |
| LC Column/MS Platform | pHILIC/QExactive | pHILIC/QExactive | pHILIC/QExactive |
| LC-MS Run Length (min) | 26 | 26 | 46 |
| Date Analysed | 2018 | 2016 | 2018 |
| Healthy Controls | 10 | 7 | 20 |
| Infected Patients | 10 | 7 | 20 |
| SRM Sets | 3 | 3 | 3 |
| MS2 data | Yes | Yes | Yes |

Information about each LC-MS experiment including the disease studied, type of study, number of controls and patients, LC-MS platform used, date when the samples from the LC-MS experiment were run and availability of fragmentation (MS2) data are presented.

### LC-MS platform

The experiments were performed at different time points (Table 1) using the same LC-MS platform: Thermo Orbitrap QExactive (Thermo Fisher Scientific) mass spectrometer coupled with a Dionex UltiMate 3000 RSLC system (Thermo Fisher Scientific, Hemel Hempstead, UK) using a ZIC-pHILIC column. While the same flow rate was used for all three datasets, the length of the run differed for DVL, which lasted longer than the other two datasets.

### Tandem mass spectrometry data

Fragmentation (MS2) of the pooled samples within each experiment was performed using higher energy C-trap dissociation (HCD) at a normalised collision energy (NCE) of 60. These were analysed with TopN data-dependent acquisition (DDA) fragmentation strategies, resulting in MS2 spectra only for some of the observed ions.

### Standard reference mixtures

As part of the quality control across each experiment, three sets of SRMs which include compounds that cover a broad range of metabolic pathways such as amino acid metabolism, central carbon metabolism and nucleotide metabolism were run twice, before and after the cohort of samples was run. Detailed information about the SRM compounds from both the +ve and -ve electrospray ionisation (ESI) mode is included in the Supporting Information (S1 Table).

## Study workflow

The data analysis process is outlined in Fig 1.

**Fig 1.**
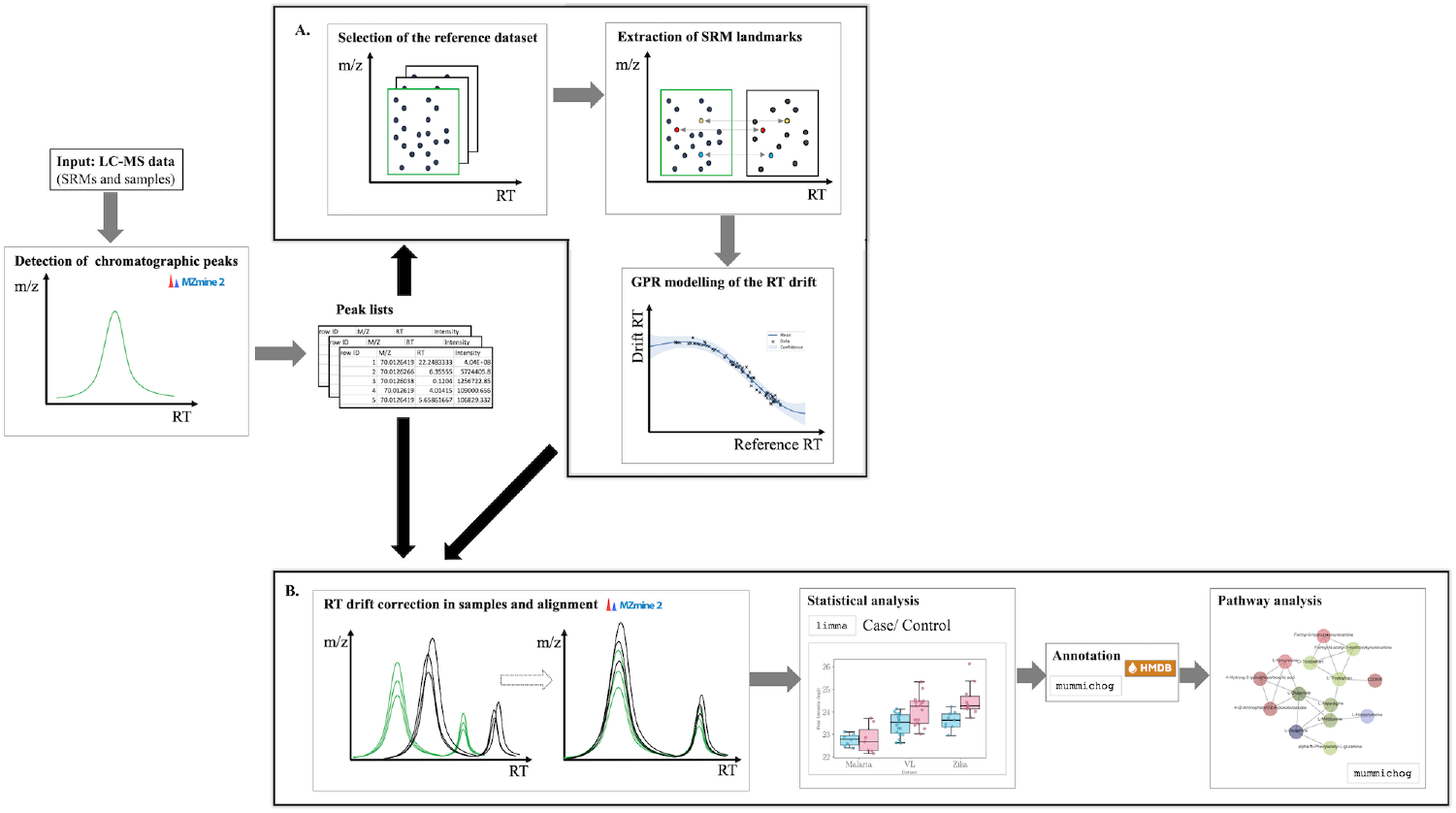
Diagram representing the study workflow. Peaks are detected from input LC-MS data including the SRMs and samples using MZmine2 and peakset lists containing ion information (m/z, RT, intensity) are obtained. A. SRM analysis. A reference dataset is selected and the RT drift in the other datasets is calculated and modelled using GP regression. B. Samples analysis. Based on the GPR models obtained for each dataset the RT is corrected in each peakset list and alignment is done using MZmine2. Afterwards, statistical analysis focused on the intensity differences between the control and infected samples is performed using limma R package. This is followed by annotation and pathway analysis using mummichog.

### Peak detection

The processing of the spectral data begins with the detection of the chromatographic peaks from the input SRM and samples LC-MS data. Peak detection was performed using wavelet transform from MZmine2 v2.40.1 [16] in batch mode.

### SRMs analysis

First, the reference dataset, D_*ref*_ = D_*Z*_, was randomly selected out of the two datasets with the shorter run length. Next, a profile was created for each dataset characterised by (m/z, RT) of the extracted SRM compounds from the peak lists obtained after the SRM peak detection process. The profiles of the non-reference datasets, D_*M*_ and D_*V L*_, were then mapped to the reference dataset profile and the RT drift between each of the non-reference datasets and the reference dataset was determined and modelled using GP regression.

### GPR modelling of the RT drift

For each non-reference dataset, the RTs from the (m/z, RT) profile created (input observations) were regressed against their respective RT drift from the reference dataset values (observed values). In order to obtain a closer fit to the data but still maintain the variability, SRM metabolite outliers were removed based on their RT drift from the reference profile using a z-score cutoff value of 2. For implementing the Gaussian Process models, the GPy python package, version 1.9.9 was used [17]. Model hyperparameter optimisation was done using multiple restarts (n=10) with the GPy optimiser to avoid local minima. To determine which covariance function aids in fitting the data best in each case, cross-validation was performed, by stratifying and splitting the SRM data in half for training and testing the model. The prediction accuracy score, mean absolute error (MAE) and mean squared error (MSE) were calculated in each case.

### GPR corrected data

The GPR corrected RT times were obtained by adding the GPR predicted variables (posterior mean) to the initial RT values of each peakset from the non-reference dataset.

### Detection of alignment RT window parameter

The alignment of the peak lists was performed using the JoinAligner module from MZmine2. The optimal RTWindow parameter was determined by aligning the SRM peak lists RTWindow values ranging from 0.01 min to 2 min. For each of the three SRM sets the total number of peaks aligned and the total number of SRM metabolites that align for each RTWindow value across the datasets is calculated. In order to determine the optimal RTWindow for all datasets, the value for which the alignment results in the lowest total number of peaks aligned and highest number of SRM metabolites is chosen for further analysis.

### Sample analysis

The GPR models obtained in the previous stage were applied to each sample peak list by correcting the RT values for each peak as detailed above. The RT corrected peak lists were then aligned using the RTWindow value previously obtained. The final list of aligned peaksets was first processed by filtering out the peaksets based on the percentage of missing values from each dataset. An arbitrary cut-off value of 50% was used. Data imputation using K-nearest neighbours method (k=3) was performed solely for visualisation purposes [18].

### Statistical analysis

The statistical analysis focused on the intensity differences between the sample peak lists belonging to the control and infected groups from all three datasets. The intensities were log2 normalised and modelled using linear regression included in the limma R package, where blocking was used to adjust for the intensity variability between the different datasets. The output of this analysis is a list containing all the peaksets and their respective p-value, Benjamini-Hochberg (BH) adjusted p-value and logarithmic fold change (logFC) between the two conditions. The formula used for the linear regression is given below.

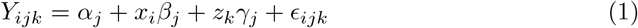

Where: *Y*_*ijk*_= response variable (intensity level of metabolite j in condition i and dataset k), *α*_*j*_ = intercept for metabolite j, *x*_*i*_= first predictor variable: condition (infected/ control), *β*_*j*_= estimated difference for metabolite j for each condition, *z*_*k*_= second predictor variable: dataset, *γ*_*j*_ = the dataset effect for metabolite j, *ϵ*_*ijk*_ = error stochastic component, within group variation.

### Feature annotation and pathway analysis

The MS2 spectra from each dataset was aligned to the final filtered peaksets list. A profile characterized by (m/z, RT, ms2spec) was created for the possible adducts/fragments for each peakset (S2 Table) and several methods were used for feature annotation. First, annotation was performed using the SRMs information by mapping the peakset profile against the SRMs (m/z, RT) profiles. Next, they were mapped against metabolite information extracted from the Human Metabolome Database (HMDB). Where one or more spectra was aligned to a peak, the attached spectrum/spectra was compared against experimental LC-MS spectra from HMDB and the match with the highest cosine similarity score was used to annotate the peak. For pathway and activity network analysis mummichog version 2.3.3. was used [19].

### Code

All of the analysis was performed in python programming language (https://github.com/anamaria-uofg/mma). The information about any given peak was stored in an object (peakinfo) with the following attributes: id, m/z, RT, p-val, t-val, logFC, mummichog annotation, mummichog pathway, mummichog kegg id, std annotation, std kegg id, spectra, adducts, best ms2 match adduct, ms2 annotation, ms2 kegg id, intensities (from each sample).

### Methodology evaluation

The peakset lists with no GPR correction were aligned and the same workflow was applied to their analysis. The results obtained were compared with the results of the GPR modified data. Also, the datasets were individually analysed and their filtered peakset lists were then intersected to determine whether any commonality can be found in this way. Additionally, we evaluated the alignment process using the available MS2 data. If two compounds with similar m/z and RT break down into the same fragments (during LC-MS analysis), then it is highly likely they represent the same compound. Therefore, if a peakset has multiple highly similar MS2 spectra from different datasets, it is likely that the peaks were aligned correctly. In order to measure the similarity between two MS2 spectra, cosine similarity score implemented in mass-spec-utils was used [20]. In order to evaluate the alignment process using MS2 data the spectral similarity between the MS2 profiles was computed when more than one spectrum was aligned to one peakset. For each of the spectral similarity scores (good spectral similarity score), the mean of the corresponding distribution of random spectral similarity scores (bad spectral similarity score) was calculated, which were obtained from spectra of peaksets with similar m/z, but different RT.

## Results

### SRM analysis

#### Correlation between the RT drift and other variables

Correlation between the RT drift and the characteristics of the dataset profiles was checked. For all datasets the highest correlation was found with the RT from each dataset profile. Based on this information, the training (70%) and test (30%) data were split and stratified in 4 equal length bins based on the RT. Following cross-validation, the kernel with the highest accuracy and lowest MAE was chosen. The final model was fitted using the selected kernel and optimised using multi-start in order to deal with possible bad local minimum.

### GPR modelling of the RT drift

Several kernels, among which RBF, neural network and cosine kernels, were tested to determine which ones best fit the data (Fig 2). Composite kernels were also tested on the data. Composite kernels refer to multiple kernels combined either by addition or multiplication.

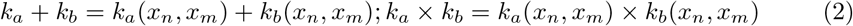

**Fig 2.**
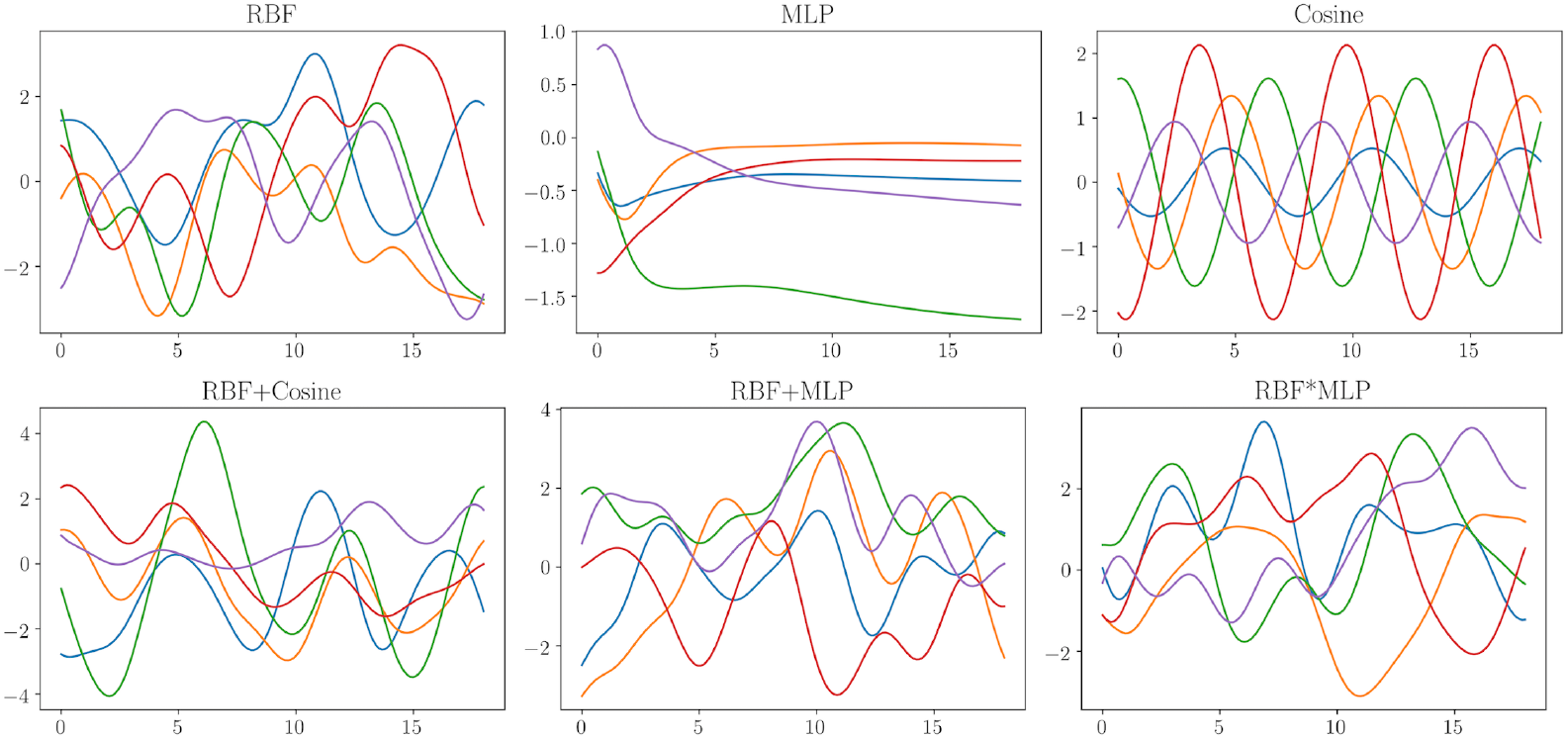
Sampling from multivariate Gaussian distribution with composite kernels. Sampling 5 times from a multivariate Gaussian distribution () with mean 0 and covariance. Covariance functions included were RBF, MLP and Cosine and composite kernels obtained either though addition (RBF+MLP, RBF+Cosine) or multiplication (RBF*MLP).

For D_*M*_ the mean retention time drift in comparison to the reference dataset was 19.85 s and the highest retention time drift between two peaks belonging to the same ion was 319.74 s. Following cross-validation an RBF kernel was selected as the best for fitting the RT drift in D_*M*_ with an accuracy of 0.99, MAE = 0.06 and MSE = 0.03. When fitted to the whole data (except the outliers) the final model had an accuracy score 0.93, MAE=0.14 and MSE = 0.46. The two hyperparameters of the RBF function, i.e. variance and lengthscale, were optimised to 0.07 and 6.84 respectively (Gaussian noise variance = 0.01) (Fig 3).

**Fig 3.**
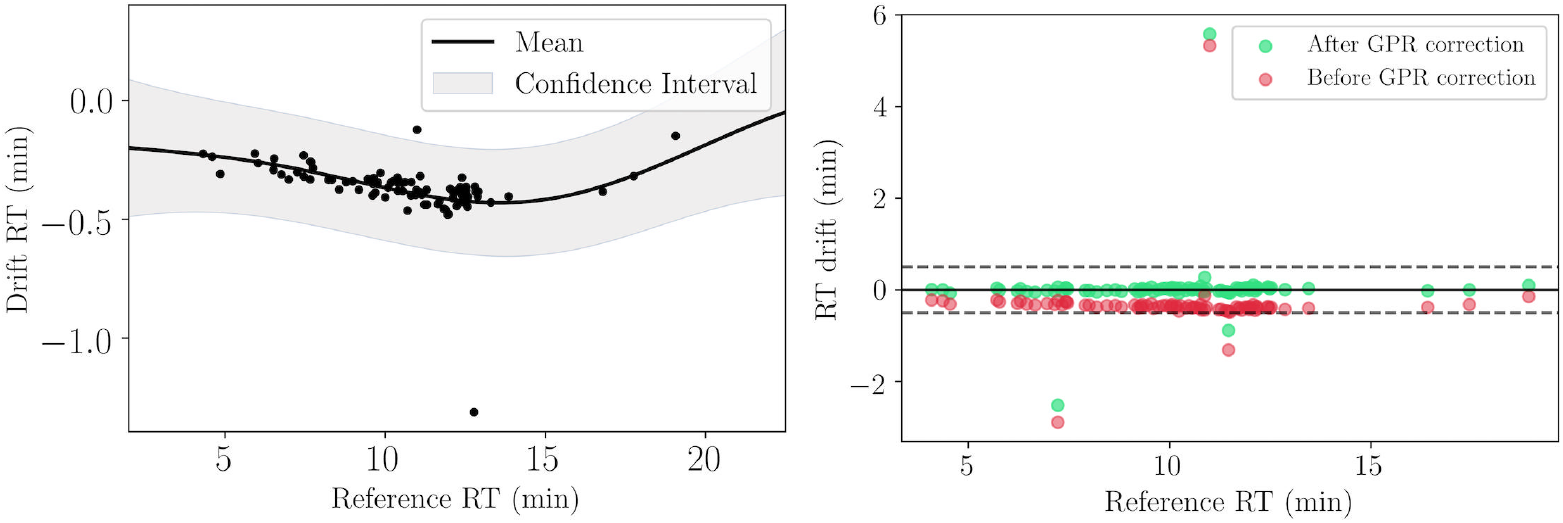
Modelling the drift in D_*M*_ using an RBF kernel. The graph on the left: Mean and 95% posterior confidence with optimised hyperparameters: RBF variance = 0.07, RBF lengthscale = 6.84. The plot on the right illustrates the RT drift in D_*M*_ before and after correction of the retention times using the GPR model.

For D_*V L*_ the mean retention time drift was 112.03 s and the maximum retention time drift was 252.54 s. For the drift in D_*V L*_ a composite kernel RBF+MLP was chosen for fitting the data with an accuracy score of 0.995, MAE=0.14 and MSE = 0.03 (Fig 4). When fitted to the whole data (except the outliers) the final model had an accuracy score 0.89, MAE=0.27 and MSE = 0.74. The hyperparameters after 10 optimisation restarts: MLP variance = 3.99, MLP weight variance = 2.02e7, MLP bias variance = 5.56e-309, RBF variance = 7.44, RBF lengthscale = 9.41 (Gaussian noise variance = 0.1).

**Fig 4.**
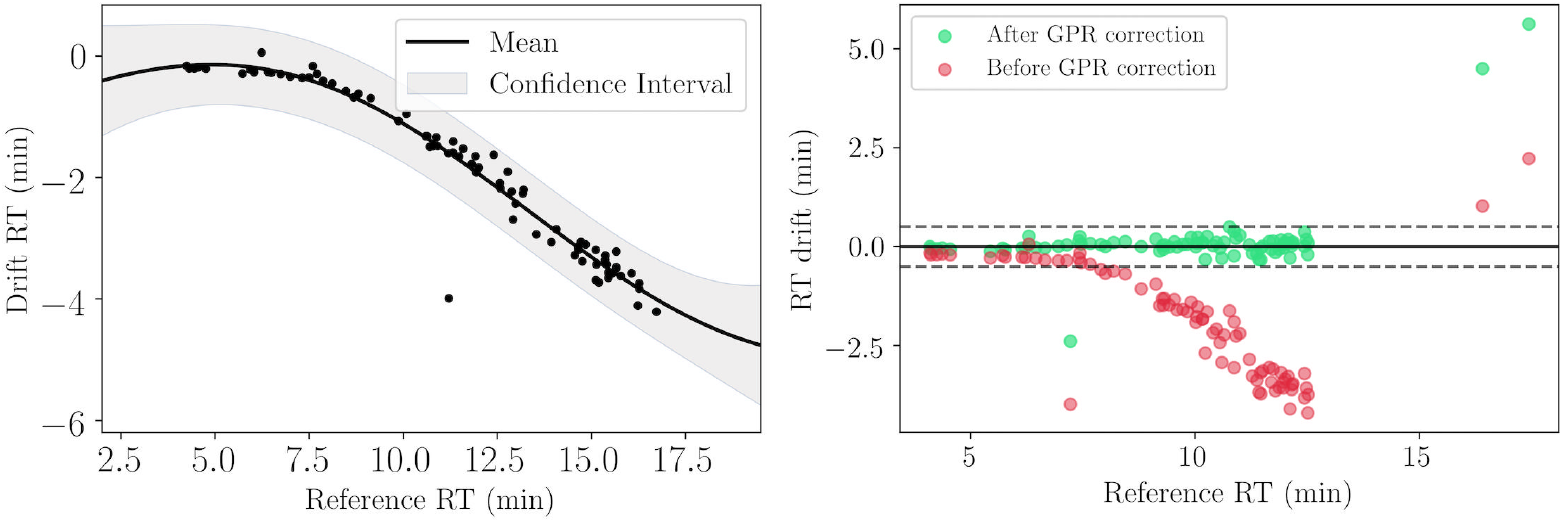
Modelling the drift in D_*V L*_ using a composite RBF + MLP kernel. The graph on the left: Mean and 95% posterior confidence with optimised hyperparameters: MLP variance = 3.99, MLP weight variance = 2.02e7, MLP bias variance = 5.56e-309, RBF variance = 7.44, RBF lengthscale = 9.41. The plot on the right illustrates the RT drift in D_*V L*_ before and after correction of the retention times using the GPR model.

### RTWindow value choice for JoinAligner module

For D_*M*_ 94.8% of the maximum number of SRM metabolites in common with the reference dataset are aligned at RTWindow = 0.25 min after RT drift correction, as opposed to 19.48% before drift correction. Whereas, for D_*V L*_ the maximum number of metabolites which are aligned after drift correction are obtained at RTWindow = 0.5 min for which 92.67% of the metabolites align, as opposed to 17.64% before correction. At a RTWindow value of 0.25 min 83.82% of the metabolites align (S1 Fig). Based on these results, the optimal RTWindow parameter for further alignment of the samples was selected, i.e. RTWindow = 0.5 min. After correction, the number of total peaksets obtained following alignment also decreases, signifying better aligned peaks. Choosing the correct parameter for the RT window is important, because if the window is too small, then false negatives are introduced and, vice-versa, if the window is too big, then false positives are introduced.

## Methodology evaluation

### GPR corrected data vs original data

Results obtained for the +ve mode data are presented. In total, 625 peaksets remained after filtering the GPR corrected data, as opposed to 344 peaksets in the original data. Most of the peaks which are aligned only in the GPR modified data have the retention time in the range of [7,13] minutes (Table 3).

**Table 2.** Correlation between the RT drift and (m/z, RT) of each dataset profile.

|  |  | D<sub>Z</sub> m/z | D<sub>Z</sub> RT | D<sub>M</sub> m/z | D<sub>M</sub> RT | D<sub>VL</sub> m/z | D<sub>VL</sub> RT |
| --- | --- | --- | --- | --- | --- | --- | --- |
| RT Drift | D<sub>Z</sub>-D<sub>M</sub> | -0.01 | -0.04 | -0.01 | 0.22 | / | / |
| RT Drift | D<sub>Z</sub>-D<sub>VL</sub> | -0.08 | 0.55 | / | / | -0.08 | 0.796 |

**Table 3.** Results obtained from the unmodified and GPR modified aligned data.

|  | Without GPR | With GPR |
| --- | --- | --- |
| Total number of aligned peaksets | 38,197 | 37,220 |
| Total number of filtered peaksets | 344 | 625 |
| Significantly modified peakset intensity values | 176 | 207 |

### Individual datasets alignment

For D_*Z*_ 2305 peaksets remain after filtering, out of which 3 are significant. For D_*V L*_ 2392 peaksets remain after filtering, out of which 775 are significant. For D_*Z*_ 2152 peaksets remain after filtering, out of which 5 are significantly different. When intersecting the significantly different peaksets, there is no peakset found to be in common between all three datasets. Therefore, aligning the peaksets together increases the number of samples, increasing statistical significance.

### MS2 data

Due to the fragmentation strategy employed in each experiment, MS2 spectra were available only for a small percentage of the data. From the filtered peaksets there were 217 peaksets with MS2 spectra aligned, out of which 141 had an MS2 spectrum from one dataset attached to it, 65 peaks had MS2 spectra from 2 datasets attached and 11 peaks had MS2 spectra from all 3 datasets. The majority of peaksets were fragmented only in one dataset. Out of the 141 peaksets with an MS2 spectrum from one dataset, 18.4% peaksets had MS2 data only from D_*M*_, 45.4% from D_*Z*_ and 36.2% from D_*V L*_. Based on Fig 5, the bad spectral similarity scores attached to peaksets with the same m/z mainly have a lower similarity score than the good scores (87%), which in general shows that the alignment worked.

**Fig 5.**
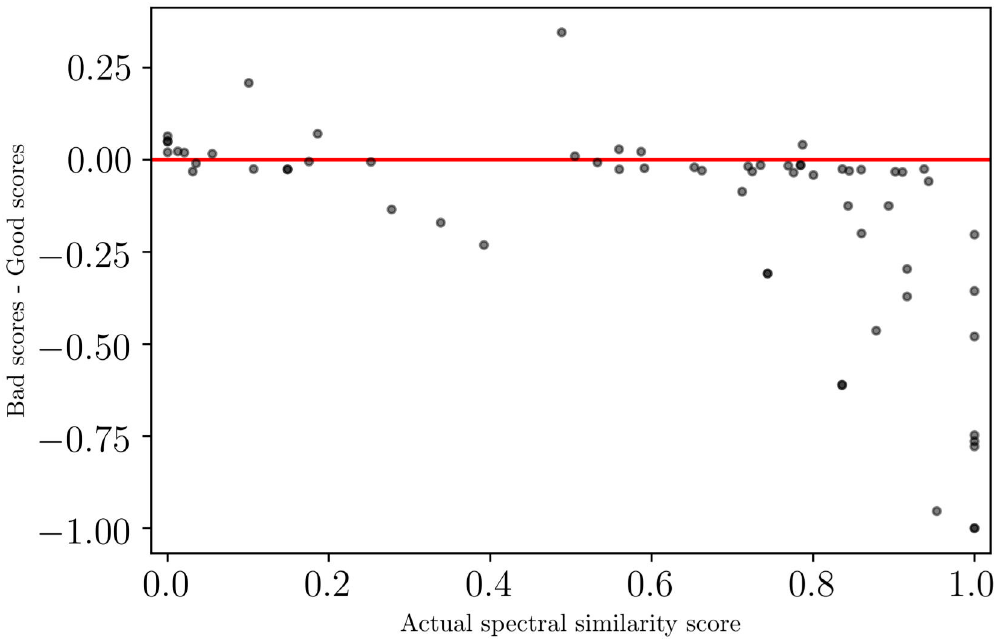
Spectral similarity scores of each peakset plotted against the difference between a random spectral similarity score and the actual spectral similarity score. The points below the red line represent the peaksets for which the actual spectral similarity score is higher than the score from randomly matched spectra with similar m/z.

## Sample analysis

### Sample alignment

For the sample alignment, the -ve ESI mode data was processed in the same way as the +ve mode data which was presented above. The JoinAligner module was run with RTWindow = 0.5 min in order to align all 74 samples across the three datasets. Following alignment there were 37220 peaksets in +ve mode and 24729 in -ve mode. After filtering out the peaksets with more than 50% values missing in any one dataset, only 1.68% of the total number of peaksets, i.e. 625, remained. A similar percentage was obtained in the case of the negative mode data where 1.85% (459) of the total number of peaksets remained. The differential expression analysis resulted in 207 significantly different (BH adjusted p-value*<*0.05) features and 159 features in the positive and negative mode, respectively.

### Overview of significantly modified compounds

The significant features with putative annotations either by standard matching, MS2 profile matching or mummichog analysis, that present common trends based on the logFC values between the two conditions (disease vs control) from each individual dataset are presented in Fig 6, 7, 8. Based on this, a general trend over the three datasets was established. For the + ESI mode data, 30 peaksets presented a general upward trend (all datasets have logFC*>*0) out of which 11 were statistically significant and 150 peaksets presented a general downward trend out of which 69 were significant. For the - ESI mode data, 46 peaksets presented a general upward trend out of which 25 were statistically significant and 115 peaksets presented a general downward trend out of which 74 were significant.

**Fig 6.**
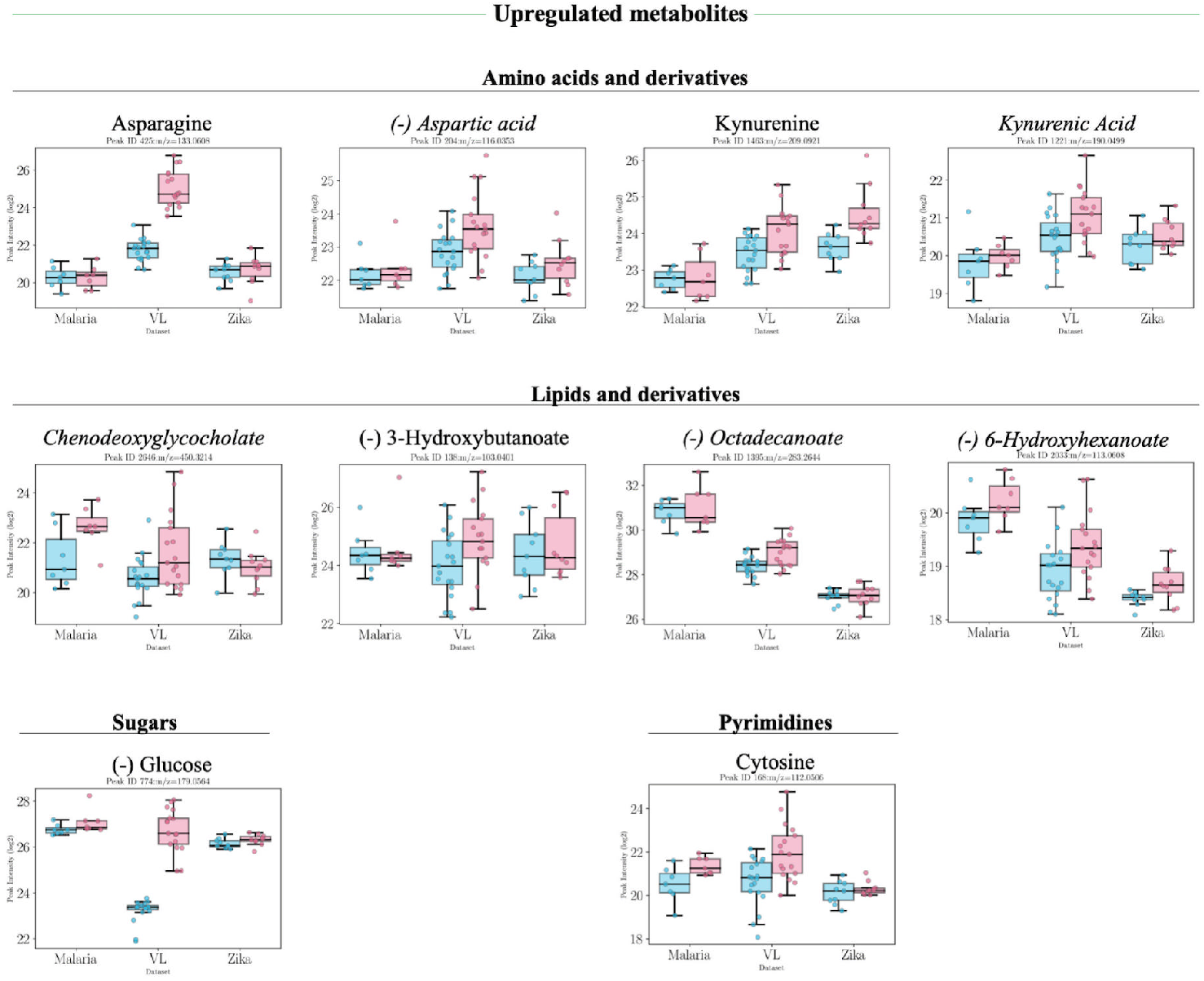
Boxplots of putatively annotated compounds for both conditions in each dataset. Overview of putatively annotated metabolites which are statistically significant (p-val*<*0.05) and present a general upward trend in all three datasets, i.e higher intensities in infected patients. Values from both positive and negative (-) ionisation mode are presented from left to right in ascending order of their p-value. Metabolites in italic font are only annotated using mummichog.

**Fig 7.**
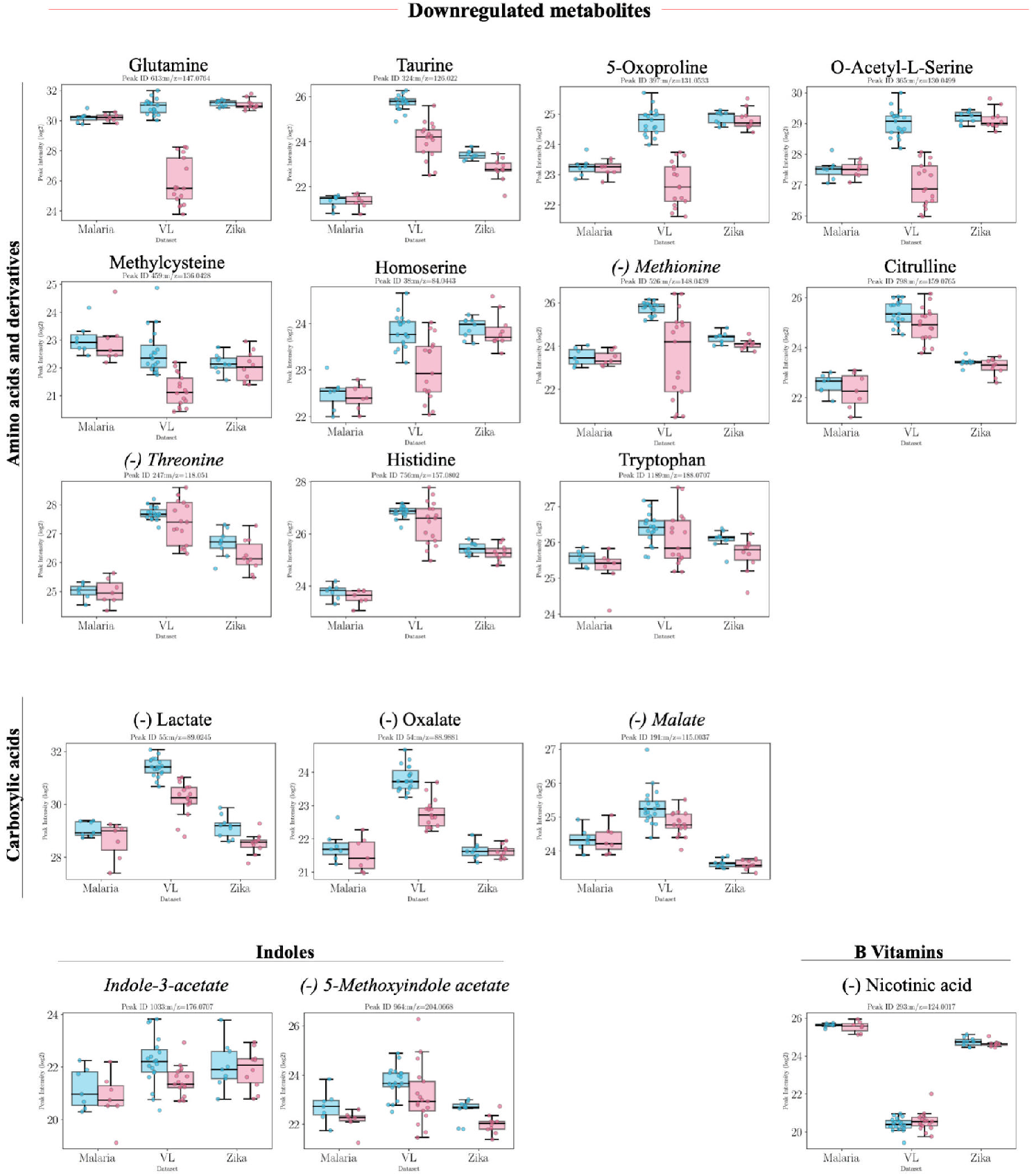
Boxplots of putatively annotated compounds for both conditions in each dataset. Overview of putatively annotated metabolites which are statistically significant (p-val*<*0.05) and present a general downward trend in all three datasets, i.e. lower intensities in infected patients (with p-value*<*0.05). Values from both positive and negative (-) ionisation mode are presented from left to right in descending order of their p-value in each group. Metabolites in italic font are only annotated using mummichog.

**Fig 8.**
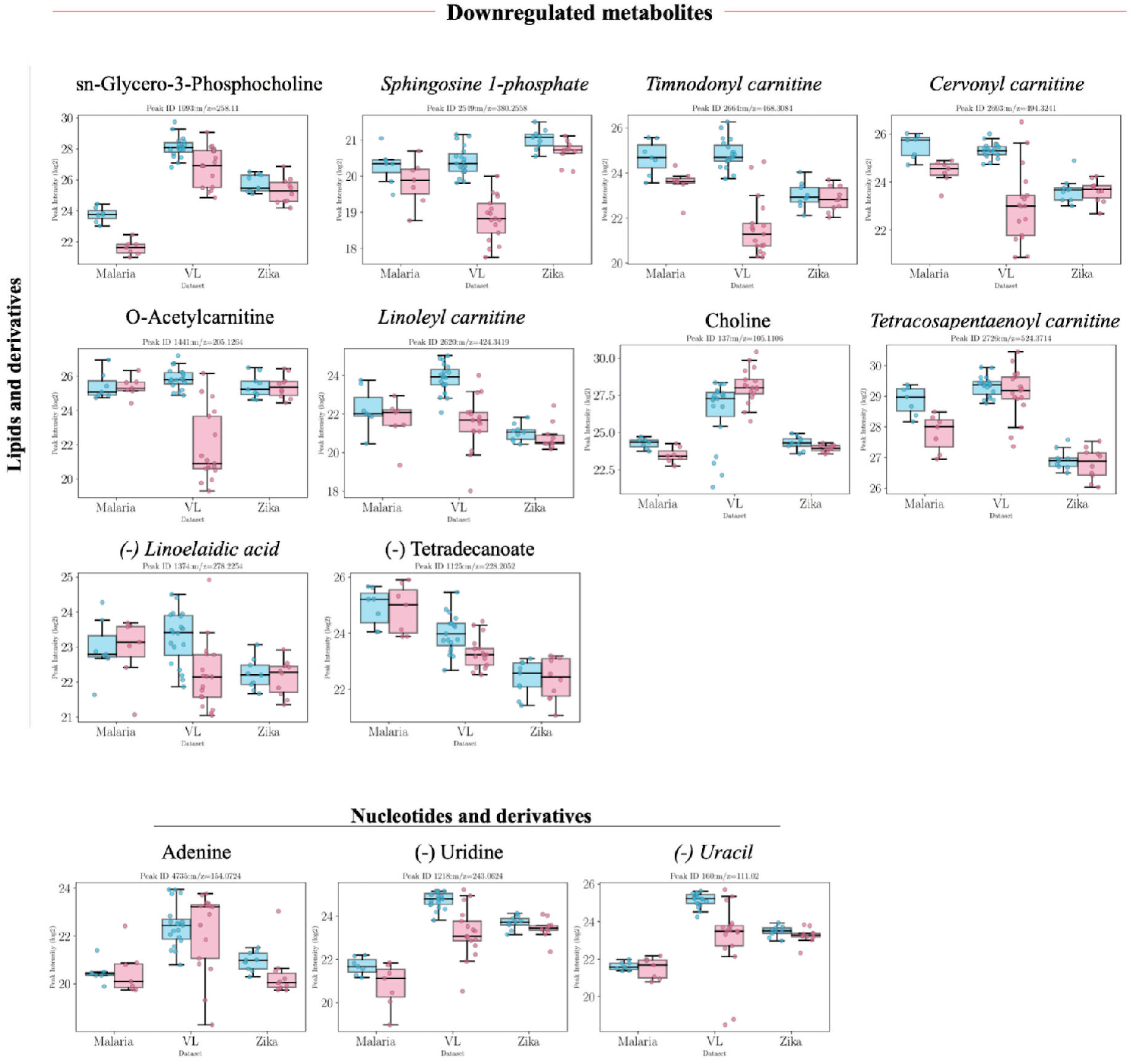
Boxplots of putatively annotated compounds for both conditions in each dataset. Overview of putatively annotated metabolites which are statistically significant (p-val*<*0.05) and present a general downward trend in all three datasets, i.e. lower intensities in infected patients (with p-value*<*0.05). Values from both positive and negative (-) ionisation mode are presented from left to right in descending order of their p-value in each group. Metabolites in italic font are only annotated using mummichog.

A table containing the results following the statistical analysis with annotation is included in the Supporting information (S2 Table). A large proportion of the features elute in clusters of similar retention time, indicating that they might be related, e.g. fragments or adducts of the parent ion. The list of ions used to calculate the possible adducts/fragments of each peak are included in the Supporting Information (S3 Table). Common adducts include gain of C^13^, S^34^, H2O, Na or K and common fragmentation patterns include loss of CO, H_2_O, HCOOH, NH_3_, C_3_H_4_O_2_, H_4_O_2_.

### Pathway analysis results

Following mummichog pathway analysis for both the positive and negative ionisation modes 20 KEGG pathways were found to be statistically significant. The pathway analysis revealed a significant impact of the studied infectious diseases on nitrogen metabolism with tryptophan metabolism predominant (Table 4). Following the modular analysis, the activity network for the + ESI mode is also centered around tryptophan metabolism, specifically the kynurenine pathway, which is discussed in more detail in the next section (Fig S3). For the - ESI mode data the activity network also involves tryptophan metabolites and, additionally, citric acid cycle metabolites.

**Table 4.** Significantly altered metabolic pathways (p-val¡0.05) following mummichog analysis of the negative and positive ionisation mode data.

| KEGG Pathway | p-value | ESI mode |
| --- | --- | --- |
| Nitrogen metabolism | 0.000672 | + |
| Tryptophan metabolism | 0.00084 | + |
| Alanine and Aspartate Metabolism | 0.002017 | + |
| Pyruvate Metabolism | 0.00521 | - |
| Vitamin B3 (nicotinate and nicotinamide) metabolism | 0.005546 | + |
| Pyrimidine metabolism | 0.007226 | + |
| Glycolysis and Gluconeogenesis | 0.008151 | - |
| Carnitine shuttle | 0.009243 | + |
| Glycerophospholipid metabolism | 0.013024 | - |
| Glycosphingolipid metabolism | 0.013024 | - |
| Methionine and cysteine metabolism | 0.015965 | + |
| Aminosugars metabolism | 0.022939 | + |
| Purine metabolism | 0.022939 | + |
| Bile acid biosynthesis | 0.031846 | + |
| Fatty Acid Metabolism | 0.031846 | - |
| Putative anti-Inflammatory metabolites formation from EPA | 0.031846 | + |
| Androgen and estrogen biosynthesis and metabolism | 0.03924 | - |
| C21-steroid hormone biosynthesis and metabolism | 0.03924 | - |
| Vitamin B1 (thiamin) metabolism | 0.03924 | - |
| Glutathione Metabolism | 0.04445 | + |

### Tryptophan metabolism

Focused analysis on the tryptophan metabolic pathways represented in Fig 9 revealed significant decreases in all three datasets of tryptophan and tryptophan derivatives such as indoleacetic acid and methoxyindole acetate. Methyl indole acetate and formyl-N-acetyl-5-methoxy kyunernamine were also significantly decreased in the infected group with the exception of the Malaria dataset where the logFC value was slightly higher (Fig 9). In contrast, the kynurenine branch of tryptophan metabolism suggests an increased activation as kynurenine and kynurenic acid are present at higher levels in infected patients in all three datasets. 3-Hydroxyanthranilate was in general higher as well with the exception of the Malaria dataset were logFC value was slightly lower. Anthranilate levels were also reduced in infected patients across all datasets, although not reaching statistical significance. Nicotinic acid (- ESI mode) was also found in significantly lower levels in infected patients and nicotinamide was significantly lower, although the Zika dataset had positive logFC (Fig 7). It is important to note that unless otherwise stated, annotations are putative and based on m/z alone.

**Fig 9.**
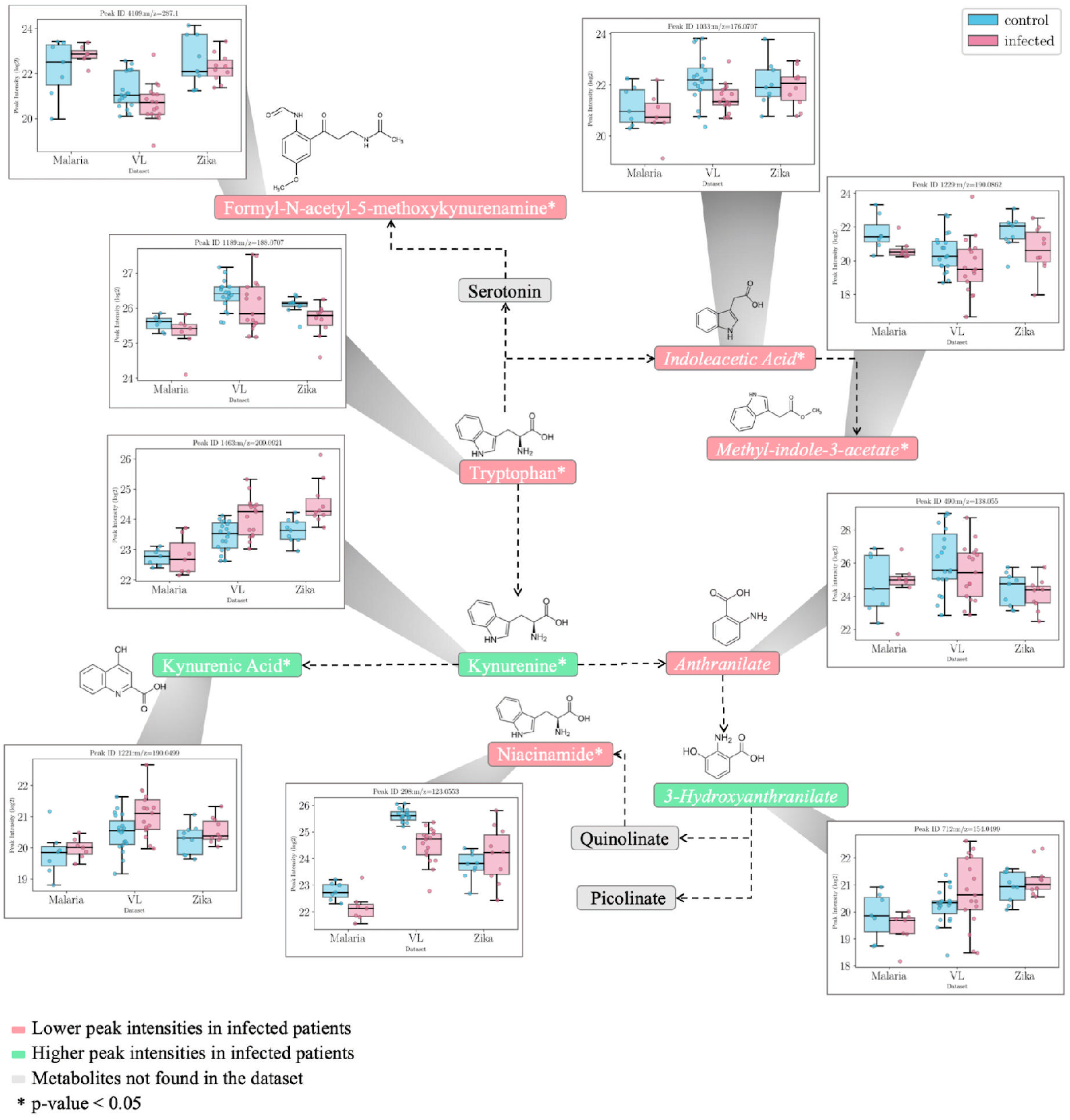
Tryptophan metabolism and the changing metabolites from each dataset. Intensity values are represented as lg2 values. The metabolites were mapped against the Kyoto Encyclopedia of Genes and Genomes (KEGG) pathway map hsa00380. Metabolites in italic font are annotated following mummichog analysis or the HMDB matching method, while the rest are annotated using the SRM metabolites information. The boxplots represent the intensities of all the samples in each condition (red = infected, blue = control) in all three datasets.

Tryptophan metabolism has previously been associated with various agents of infection [21], particularly its flow through the kynurenine pathway which produces metabolites including kynurenate and nicotinamide adenine dinucleotide (NAD+). Of particular interest is the inverse relation between kynurenine and tryptophan, as the ratio between the two is used to measure the activity of the enzyme indoleamine-2,3-dioxygenase 1 (IDO-1) [22]. IDO-1 is the rate limiting step of the tryptophan pathway and it catalyses the breakdown of tryptophan to kynurenine. IDO-1 activity is also tightly regulated by interferon gamma (IFN-*γ*) activity [23]. Similar to this, COX-2, an enzyme central to the fever process, can also be induced by IFN-*γ* [24]. The interplay between IDO-1 and COX-2 enzymes has been previously studied where inhibition of COX-2 enzyme has led to a downregulation in IDO-1 and decrease in kynurenine metabolites [25].

It could be hypothesised in this case, that the increased level in kynurenine and decreased levels of tryptophan indicate an increased IDO-1 activity, and subsequently an increased COX-2 activity. Due to the importance of these metabolites further investigation was conducted in verifying the identity of the putatively annotated metabolites using the available MS2 spectral information.

### Verifying the identity of tryptophan and kynurenine peaks using MS2 spectral information

As both the tryptophan and kynurenine peaks had spectral information from one out of the three datasets, it was used to verify the identity using the cosine similarity score with experimental LC-MS/MS spectral information from HMDB. Metabolomics spectrum resolver was also used to further check the similarity between the dataset spectrum and spectra obtained from MassBank. Similarity scores of 0.35 and 0.54, respectively, were obtained for kynurenine and 0.31 and 0.55, respectively, for the tryptophan fragment (with loss of ammonia) (Fig 10, Fig 11).

**Fig 10.**
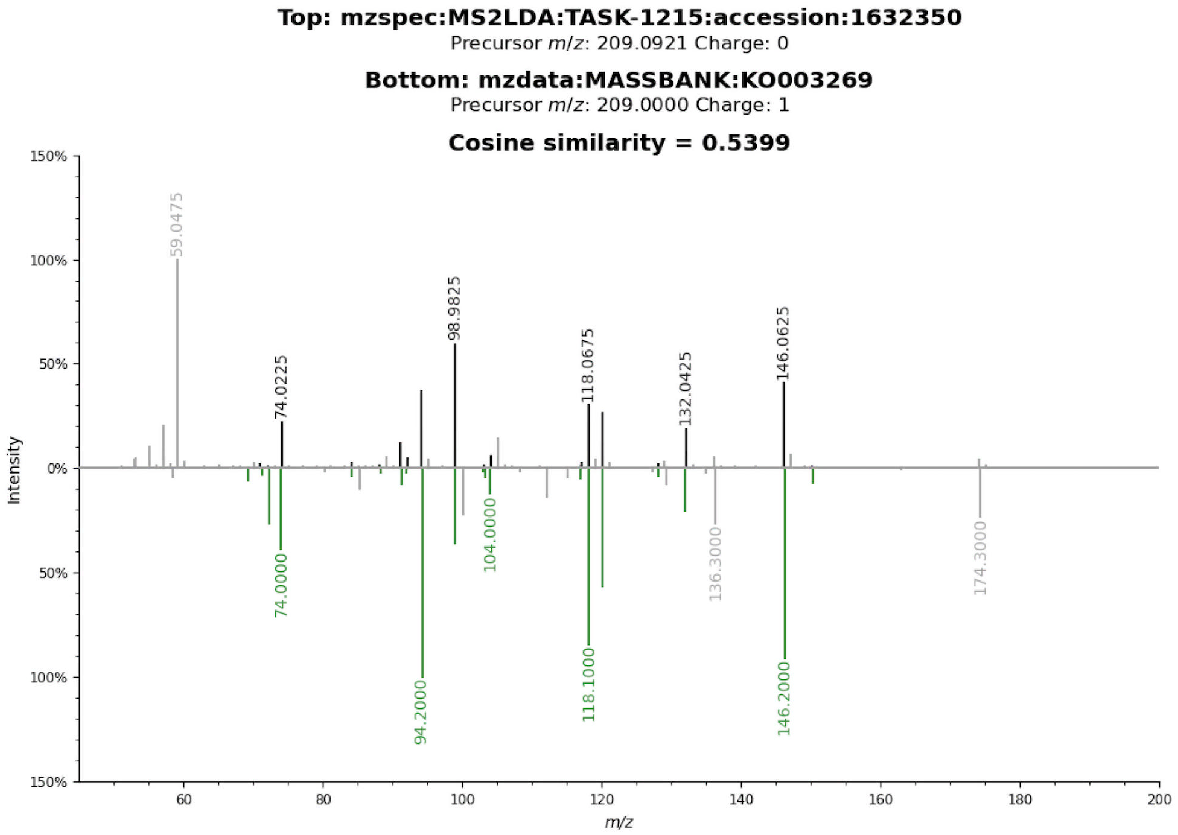
Peakset annotated as Kynurenine (M+H[1+]). using SRM matching method, mummichog and HMDB matching method. The spectrum belongs to D_*V L*_ (resolver obtained from ms2lda.org) and it was matched against experimental LC-MS MS2 information from MassBank compound KO003269 with a cosine similarity score of 0.54 (fragment tolerance =0.2) (Metabolomics spectrum resolver plot).

**Fig 11.**
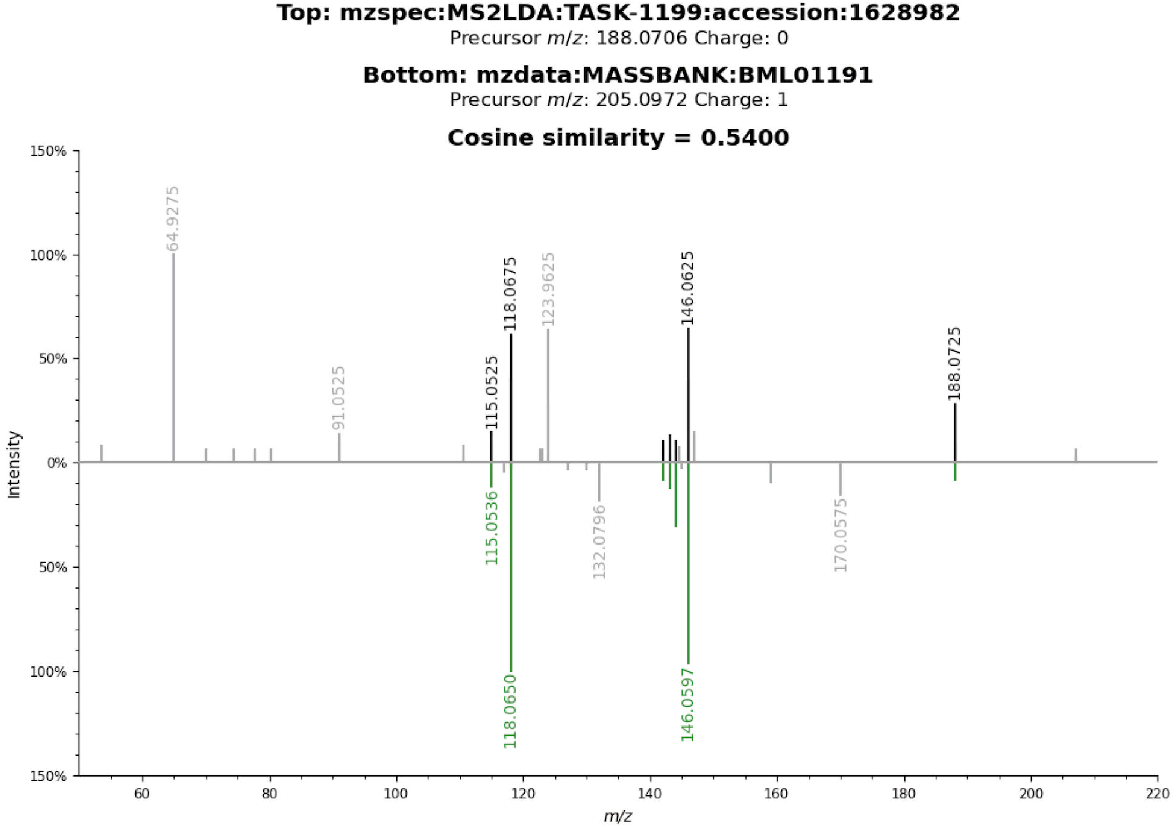
Peak annotated as L-Tryptophan fragment with loss of ammonia (M-NH3+H[1+]). using SRM matching method, mummichog and HMDB matching method. The loss of ammonia from protonated tryptophan was observed as the primary fragmentation pathway in gas-phase reactions [26]. The spectrum belongs to D_*Z*_ and it was matched against experimental LC-MS MS2 information from MassBank BML01191 compound with a cosine similarity score of 0.55 (fragment tolerance =0.2) (Metabolomics spectrum resolver plot).

### Verifying the alignment in the case of niacinamide

Both niacinamide and niacin levels are significantly lower in infected patients. The peak annotated as niacinamide in + ESI mode has MS2 spectra from two datasets and in this case the similarity between the two spectra is illustrated in Fig 12. A cosine similarity score of 1 was obtained, demonstrating that in this case the alignment between the datasets was optimal.

**Fig 12.**
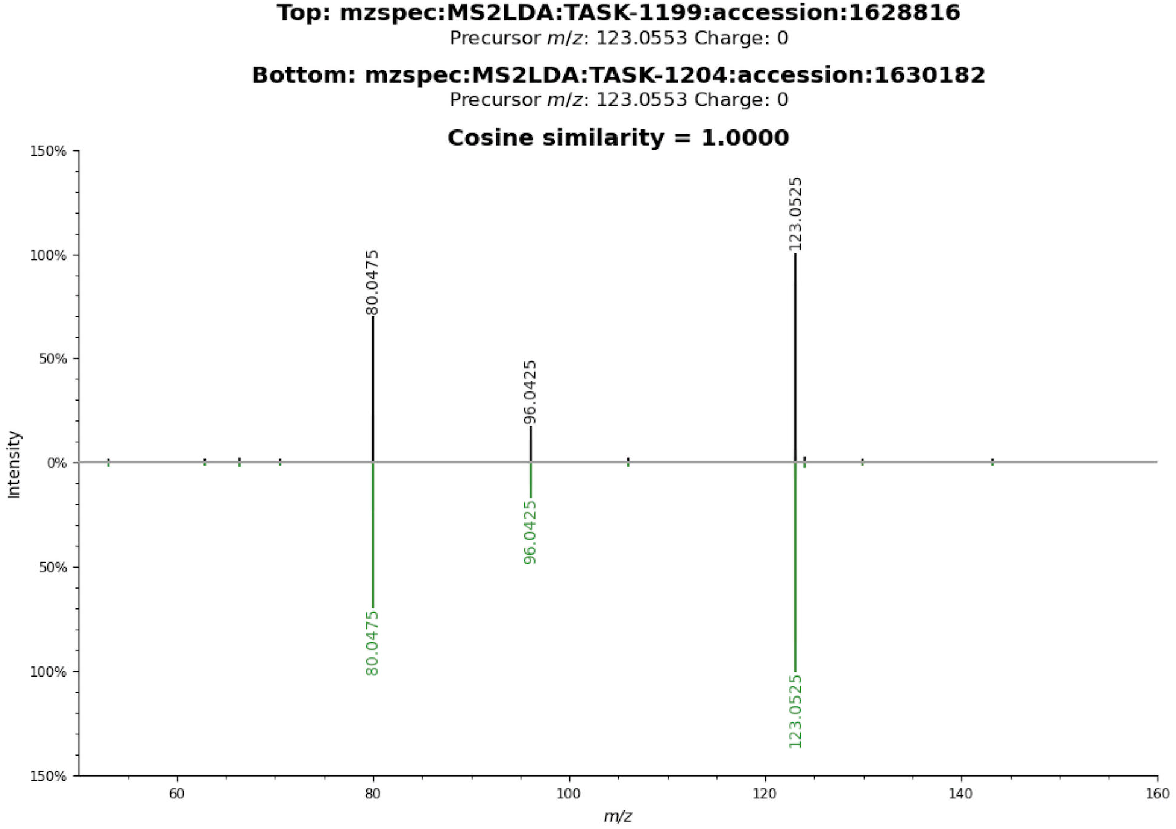
Peak annotated as Niacinamide (M+H[1+]) using SRM matching method, mummichog and HMDB matching method. Top spectrum belongs to the MS2 information obtained from D*M* and the bottom spectrum belongs to the MS2 information obtained from D_*Z*_. The cosine similarity obtained using metabolomics spectrum resolver is 1, which signifies a perfect match and also that the alignment was accurate in this case (Metabolomics spectrum resolver plot).

## Other affected metabolic pathways

### Amino acid metabolism

Other significantly affected pathways relate mostly to other amino acids including alanine, aspartate and glutamate metabolism, methionine and cysteine metabolism and glutathione metabolism (Table 4). Figure 6 indicates a clear reduction in a number of amino acids and their metabolites particularly glutamine and related metabolites. In immune cells, glutamine is converted through glutaminolysis to glutamate, aspartate and alanine by undergoing partial oxidation [27]. This could explain the decrease in glutamine and increase in aspartate in the infected group in all three datasets. Similarly, another glutamine derived metabolite, asparagine, was found to be increased in infected patients in all three datasets.

Glutamine also acts as a precursor for citrulline which plays an important role in arginine biosynthesis in the urea cycle. Citrulline levels were significantly lower in the infected group from the three datasets.

Altered glutathione metabolism accompanied by changes to metabolites associated with sulfur containing amino acids and their metabolites including methionine and cysteine metabolism could also be observed. The plasma levels of the amino acid 5-oxoproline (pyroglutamic acid) were also lower in the infected patients in comparison to healthy controls. In addition to decreased methionine, methylcysteine was also decreased in the infected patients. A precursor of cysteine, o-acetylserine was also decreased in the infected group alongside threonine and homoserine. Taurine, another metabolite derived from cysteine metabolism, was also found to have significantly lower levels in infected patients in the metadataset. The decrease in reduced thiols, may arise due to increased levels of oxidative stress in response to infection.

Other amino acids presenting lower levels in all three datasets in the infected group but not with statistical significance include: beta-alanine, proline and betaine. In contrast, several amino acids and their relatives presented overall higher levels in infected patients including carnitine, tyrosine and leucine.

### Carbon metabolism

A significant increase in glucose was also noted across datasets (Fig 6) indicating alterations in central carbon metabolism. Indeed, perturbations to the glycolytic process and citric acid cycle, in particular, were confirmed in - ESI mode data particularly with significant decreases in lactate, oxalate and malate.

### Lipid metabolism

Lipid abnormalities were also noted in the sera of infected patients, which demonstrated significant changes in the fatty acid metabolism with significant decreases in particular acylcarnitines (Fig 6). Fatty acids, hydroxybutanoate, 6-hydroxyhexanoate and octadecanoate were, on the other hand, all increased. This could be related to the previously mentioned increase in carnitine and subsequent decrease in acetylcarnitine levels which might indicate an increased activity in releasing fatty acids. Short fatty acids are known to have a protective role and play a beneficial role in reducing endothelial activation, which leads to a reduction in cytokine production and adhesion molecule expression [**?**].

Sphingolipid and glycerophospholipid metabolism were also affected with sphingosine 1-phosphate diminished in infected samples in all datasets. This might be indicative of liver damage, which is also reflected in the significantly modified levels of bile acids and taurine. Levels of taurine decrease significantly in infected patients while the bile acid chenodeoxycholate is increased in patients. Choline and its derivatives were also downregulated in infected patients, which might also be related to macrophage metabolism.

A compound with the mass of linoelaidic acid or linoleic acid was significantly lower in infected samples across all datasets. A second feature with the same mass, annotated both by SRM matching and MS2 spectral validation as linoleic acid was also decreased in infected patients, albeit without reaching statistical significance. Linoleic acid is a precursor for arachidonic acid from which prostaglandins and other bioactive eicosanoids are synthesised. In the context of fever, it could be hypothesized that an increased production of PGE2 due to the activation of COX-2 causes an increased turnover of arachidonic acid with a subsequent increased usage of linoleic acid, hence the decreased levels in serum. The LC-MS platform used here does not readily detect arachidonic acid or its products.

### Nucleotide metabolism

Pyrimidine metabolism is also affected with cytosine being significantly higher in infected patients. Viral infections are known to cause metabolic changes in host cells, including upregulation of pyrimidine nucleotide biosynthesis [28]. Uracil and its derivative, uridine, however, were found to be significantly decreased in all three datasets in the infected group. Purine metabolism is also affected, with adenine being significantly lower in infected patients

## Discussion

### Alignment of disparate LC-MS datasets

In this study we have identified a set of metabolic compounds some of which were putatively annotated which acted in a similar manner in three different LC-MS case-control datasets including infectious diseases associated with fever. The diseases were malaria, Zika virus disease and visceral leishmaniasis. The algorithm behind the integration of the three datasets was based on alignment using the raw files with corrected RT. Correspondence algorithms have been studied and variation in retention time can be classified into system variation and component level variation [9]. The system variation is general to the whole run, whereas component level variations are specific to a single analyte or a group of analytes, so cannot be modeled using monotonic functions.

Recent studies which have explored the retention time drift problem in the context of large sample sizes have led to several algorithms being developed to correct the problem. One study [29] presented a method for aligning the samples at a population scale (N=2895 human plasma samples) by correction of the non-linear retention time shift inside the raw files, in a manner similar to that developed here. In order to determine the retention shift between samples, isotope labelled standards were used to allow modelling of the shift to correct raw files for further peak detection and alignment. Our approach also corrects raw files prior to peak detection and alignment but does not require heavy isotope labelled standards. Another study in which alignment between samples from a large-scale dataset (N = 1000) is addressed [30] proposed a profile-based alignment algorithm which uses a graphical time warping method to correct the retention times for mis-aligned features.

A few studies used endogenous reference peaks to model the retention time shift between sets of samples. Li et al. [31] for example, used adjacent tandem mass spectrometry information to select endogenous reference compounds to model RT drift. A recently published study [32] also used internal reference compounds selected based on their m/z and intensity to model the RT difference between two aligned LC-MS datasets using a generalised additive model.

The advantage of aligning multiple LC-MS datasets at a retention time level over comparing previously annotated results from different datasets is that both putatively annotated and unannotated compounds that act in the same way or uniquely to each disease can be detected. Additionally, as the overall sample size is increased, by including separate but related datasets, statistical robustness of the analysis is enhanced, provided assumptions on the underlying similarity in responses of disparate datasets (for example separate pathogen related infections here) are robust. A common limitation in many LC-MS biomarker discovery studies introduced by the small sample size is thus overcome. Annotation using MS2 information is also improved, as some datasets contain MS2 spectra for peaksets which are absent in other datasets. This could be advantageous for datasets which have limited fragmentation data available, as it improves the chance of a peakset having MS2 spectra aligned, and thus the possibility for better annotation.

A limitation of our study was that the algorithm was only tested with datasets run in the same laboratory, on the same LC-MS platform and at the moment it is not known whether this method could be applied to metabolomics datasets run on different platforms. Annotation is also a limitation, as it is for metabolomics studies in general [33, 34]. It is to be noted that for some of the metabolites, the difference between the control group and the infected group was larger in one of the datasets than in the other datasets and in some cases this could have contributed to the statistical significance of the difference in the metabolites abundance in control as opposed to infected. This could be either due to the disease itself or its severity. In this case it is to be noted that DM was an intervention study and the infection was controlled and less severe which might explain relatively lower logFC in the dataset. However, for this study only the metabolites which presented fold changes going in the same direction for all datasets were presented and discussed unless specified otherwise. Using disparate diseases of matched severity might identify even more common features, but our study was constrained by the availability of particular datasets.

### Pathways affected in common by infection with separate fever-related pathogens

The algorithm was written with the intention of comparing datasets related to infection, hence we sought differences between infected patients and healthy controls from all three datasets using linear regression from limma and mummichog to determine the biological network of activity and significantly affected pathways common to the infected samples group. This study offers a route to identify commonality in the metabolic profile in infected patients affected by pathogens that cause fever. At the centre of this meta-dataset stands the relationship between kynurenine and tryptophan related metabolites.

The currently proposed molecular basis for fever involves the following. The innate immune system is activated through pathogen recognition by toll-like receptors e.g. TLR-4. This initiates the production of pyrogenic cytokines (IL-6, IL-1b, TNF-a). These pyrogenic cytokines then act on the organum vasculosus of the laminae terminalis in the hypothalamus. At the same time, PGE2 is released from hepatic Kupfer cells via the activation of cyclooxygenase-2 (the rate limiting enzyme in the synthesis of prostaglandins). PGE2 also acts on the pre-optic nucleus in the hypothalamus leading to an elevated temperature set-point. Additional negative feedback systems prevent excessive elevation of body temperature via antipyretic cytokines (IL-1RA, IL-10, TNF-a binding protein) [1].

Based on the results obtained here, a possible connection between fever and the kynurenine pathway could be explained by the interplay between IDO-1 and COX-2. An activation of IDO-1 could lead to a decreased inhibition of COX-2 which in turn could lead to an increased activation of PGE2 release. The link between the two enzymes and inflammation has been previously studied [24]. Suppression of COX-2 enzyme activity could also be decreased by the lower levels of nicotinamide in the infected group, as nicotinamide has been shown to influence the activity of COX-2 [35]. It is worth noting that serum metabolomics, as used here, detects only a faint echo of the changes that are occuring in specific cell types orchestrating inflammation in local anatomical sites associated with infection. Although PGE2 was not annotated in these datasets (PGE being difficult to detect using the platform used), linoelaidic acid and linoleic acid were putatively annotated and found to be decreased. Linoleic acid is a precursor of arachidonic acid and bioactive eicosanoids including PGE2. Decreased levels of linoleic acid in the infected group could corelate to its increased use in arachidonic acid and, thus, PGE2 production. Recent metabolomics investigations on SARS-CoV2 infection in coronavirus patients also pinpointed significant alterations in the tryptophan-kynurenine pathway [36, 37], with kynurenine levels increasing with disease severity.

Many of the other metabolites found here to have significant abundance level differences between the two groups have also been noted previously for roles in immunometabolism. Glutamine, for instance, plays a major role in the immune system, as a key energy source for immune cells and it is used at a higher rate during catabolic conditions such as sepsis or other infections [27]. Depletion of glutamine and also citrulline identified in this study could also be used as indicators of disease severity as suggested in [38]. Decreased oxoproline levels were also previously associated with a non-infectious fever-associated disease, Rheumatoid Arthritis [39]. Taurine is another amino acid which has previously been associated with inflammation and fever associated with diminished levels of the inflammatory cytokines TNF-*α* and IL-6 [40], and also related to lower body temperature when administered intracerebroventricularly [41].

## Conclusion

In conclusion, three LC-MS datasets obtained separately to investigate metabolic changes in Zika, malaria and VL infected patients were successfully aligned using fitted GP models for correcting the RT drift between them, determined by the RTs of the SRM metabolites within each dataset. Following annotation and statistical annotation, compounds changing in abundance in similar ways were found across the different infectious diseases. Common dysregulation patterns were observed in metabolic pathways related to amino acid and carboxylic acids metabolism, lipid and nucleotide metabolism with kynurenine pathway from tryptophan metabolism being identified as the most significantly changed pathway.

For more information, see Supporting information.

## Supporting information

**S1 File. Details regarding the sample collection for each dataset.**

**S1 Table. SRM metabolites information**.

**S2 Table. List of filtered peaksets**.

**S3 Table. Table with positive mode ion adducts.**

**S1 Fig. Checking the SRM alignment**.

**S2 Fig. Chromatogram of corrected RT samples.**

**S3 Fig. Mummichog activity networks**.

**S4 Fig. Linoleic acid in control vs infected samples**.

