## Supplemental Information for "Alignment of multiple metabolomics LC-MS datasets from disparate diseases to reveal fever-associated metabolites"

### Appendix

#### S1 Details regarding the sample collection for each dataset

Samples were taken from three separate LC-MS studies into different infections with pathogens associated with febrile disease. For $D_{VL}$, plasma was taken from 20 adult patients from Fuenlabrada (Madrid, Spain) diagnosed with visceral leishmaniasis at the Hospital Universitario de Fuenlabrada between January 2013 and June 2015. Blood samples were collected during the period of active disease and infection by *Leishmania infantum* was confirmed by Leishmania specific nested PCR; the presence of Leishmania-specific plasma antibodies were determined by rK39 immunochromatographic test (Inbios, USA) and indirect immunofluorecence test. Data obtained were compared with those of 20 matched healthy controls obtained from volunteers at the Blood Bank of the Hospital Universitario de Fuenlabrada [1]. For $D_Z$, patients included in the Zika virus group had presented to hospital seeking assistance for febrile symptoms confirmed as Zika virus infection by PCR (n=10). The healthy control group was composed of women attending their routine prenatal care (n=10). Samples from all participants were collected by using a red cap vacuum blood tube of 4 mL, without clot activator. After blood collection, the tubes were maintained in the rack, at room temperature for 30 minutes. After this time, samples were maintained at 4°C until centrifugation (which was performed in less than 5 hours). Centrifugation was carried out at 2000 x g for 10 minutes. Serum samples were then placed in 1.5 microtubes and stored at -80°C until metabolites extraction [2]. Regarding $D_M$, the malaria patients are described in detail in [3].

#### S1 Table. SRM metabolites information (Google sheets link)

#### S2 Table. List of filtered peaksets (Google sheets link)

#### S3 Table. Table with positive ESI mode ion adducts (Google sheets link)

[2] Botana L, Matia B, San Martin JV, Romero-Mate A, Castro A, Molina L, et al. Cellular Markers of Active Disease and Cure in Different Forms of Leishmania

Infantum-Induced Disease. *Frontiers in Cellular and Infection Microbiology*.2018;8. doi:10.3389/fcimb.2018.00381.15.

[3] Milne K, Ivens A, Reid AJ, Lotkowska ME, O'Toole A, Sankaranarayanan G,et al. Mapping Immune Variation and Var Gene Switching in Naive Hosts Infected with Plasmodium Falciparum. *eLife*. 2021;10:e62800.doi:10.7554/eLife.62800
