## Supplementary figures and images for "Alignment of multiple metabolomics LC-MS datasets from disparate diseases to reveal fever-associated metabolites"

### S1 Fig. A

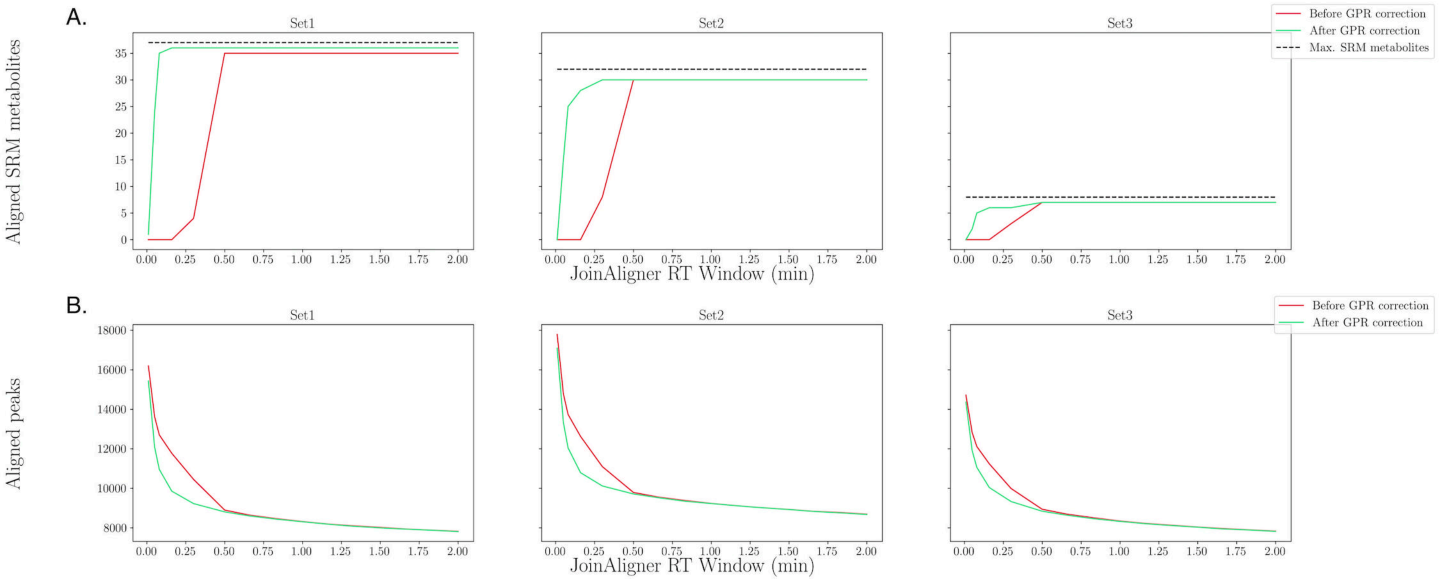

### S1 Fig. B

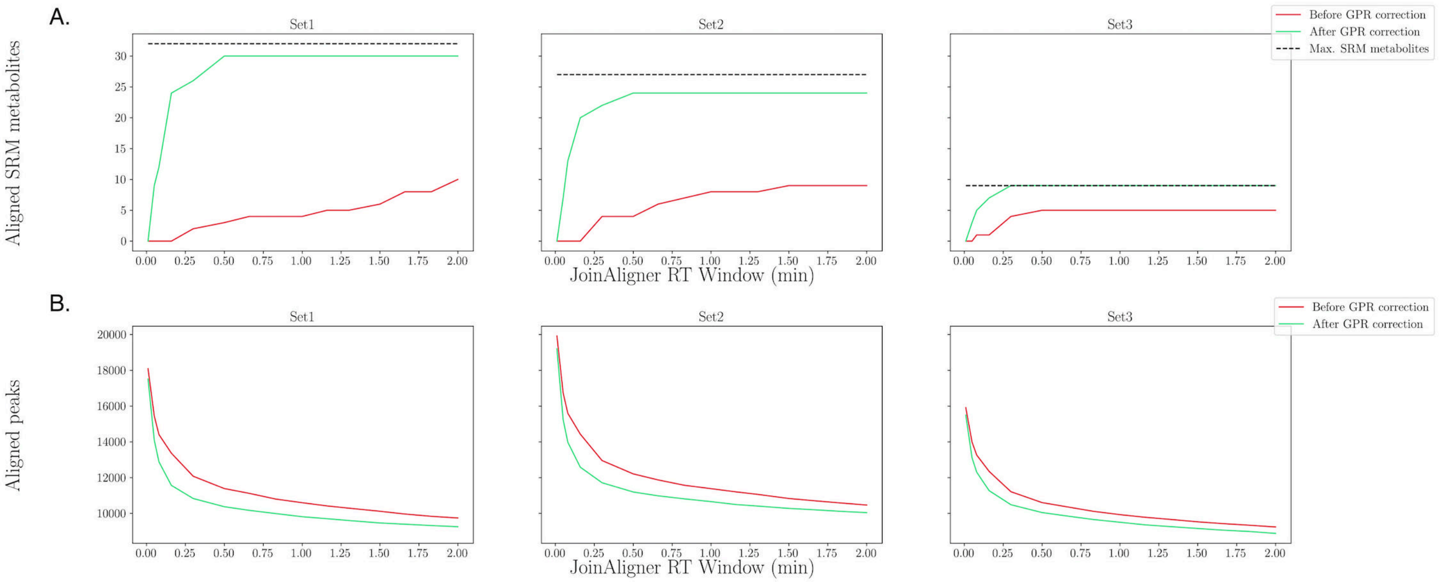

### S2 Fig.

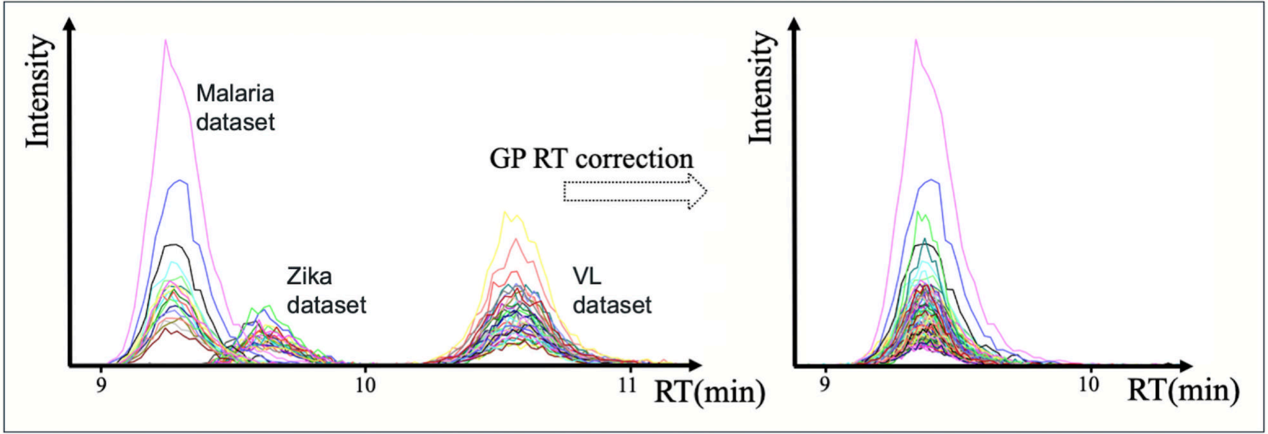

### S3 Fig.

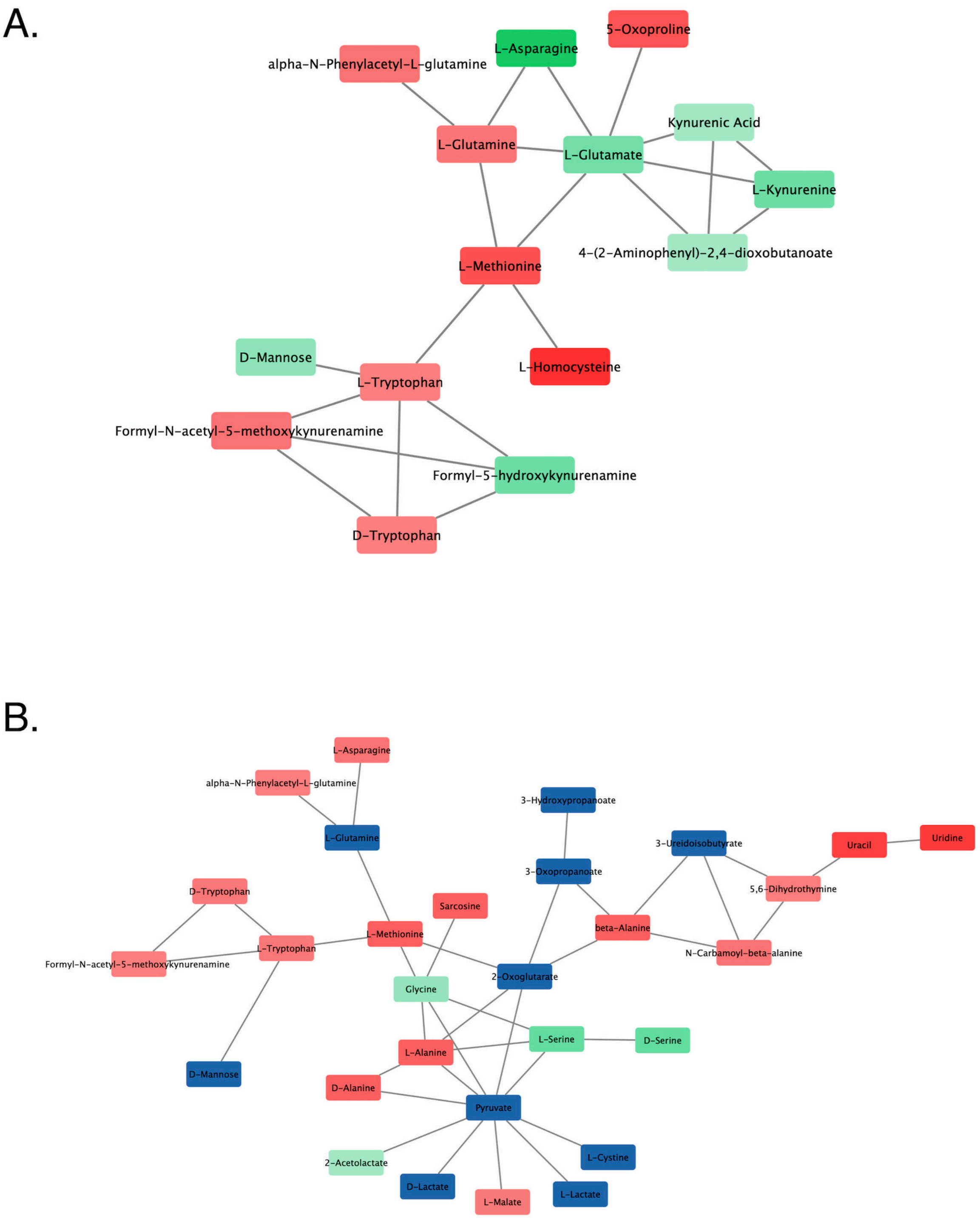

### S4 Fig.

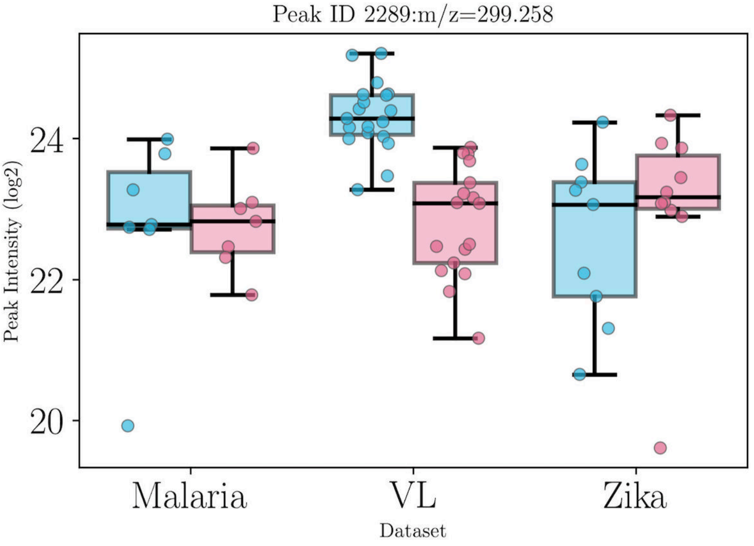

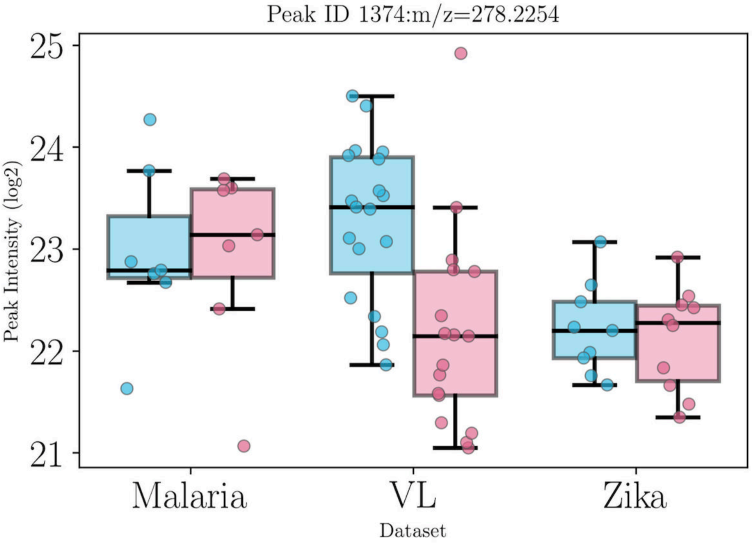
